# Novel Highly Pathogenic Avian Influenza (A)H5N1 Triple Reassortant in Argentina, 2025

**DOI:** 10.1101/2025.05.23.655175

**Authors:** Ralph E. T. Vanstreels, Martha I. Nelson, María C. Artuso, Vanina D. Marchione, Luana E. Piccini, Estefania Benedetti, Alvin Crespo-Bellido, Agostina Pierdomenico, Thorsten Wolff, Marcela Uhart, Agustina Rimondi

## Abstract

Genomic sequencing of re-emerging highly pathogenic avian influenza A(H5N1) virus detected in Argentina in February 2025 revealed novel triple-reassortant viruses containing gene segments from Eurasian H5N1 and low pathogenic viruses from South and North American lineages. These findings underscore continued evolution and diversification of clade 2.3.4.4b H5N1 in the Americas.

---

Highly pathogenic avian influenza (HPAI) was introduced to South America in 2022, by migratory birds from North America. The viruses belonged to the 2.3.4.4b clade of A(H5N1) that became widespread in Europe in 2020 and spread to North America in 2021. The trajectory of H5N1 in South America has differed from North America in several important ways. First, almost all South American H5N1 cases stem from a single introduction of H5N1 (B3.2 genotype) viruses from North America (1,2), whereas the North American H5N1 epizootic has been re-seeded by multiple independent introductions from Europe and Asia (A1-A5 genotypes) (3). Second, the South American H5N1 outbreaks were driven by a unique genotype (B3.2) that remained genetically stable as it spread across the continent and host-switched to mammals. In contrast, the H5N1 virus in North America has undergone frequent reassortment with low pathogenic avian influenza (LPAI) viruses, prompting new genotype nomenclature (B, C, D) to track the emergence of novel reassortants (3). Third, the South American H5N1 epizootic is unique for establishing mammal-to-mammal transmission in wild marine mammals (1). This transmission was likely facilitated by the H5N1 (B3.2) virus acquiring mammalian-adaptive PB2 mutations (Q591K, D701N), which then enabled its spread among mammals over extensive distances and caused recurrent mortality events in Peru, Chile, Argentina, Uruguay, and Brazil (1,2). Moreover, the presence of both PB2 mutations in the only severe human case reported in Chile in March 2023 highlights the zoonotic potential of this genotype (4). This pattern has not been observed in North America, where H5N1 has frequently spilled into terrestrial and marine mammals but only transiently, except for spread in U.S. dairy cattle (3).

Beyond the ecological devastation among coastal wildlife, in 2023 the H5N1 (B3.2) virus spread widely in birds across mainland South America, leading to poultry and wild bird outbreaks in Peru, Chile, Argentina, Uruguay, and other countries (5–8). Although in 2024 HPAI outbreaks ocurred in Brazil and Peru (9), there were no detections in Argentina from March 2024 to January 2025.

On 11 February 2025, Servicio Nacional de Sanidad y Calidad Agroalimentaria (SENASA) was notified of an outbreak in a mixed backyard flock (chickens, ducks and turkeys) in Chaco province, northern Argentina (26.3993°S, 60.6797°W). The flock experienced high mortality (33 of 81 chickens and 37 of 99 ducks died over one week). Upon inspection on the following day, approximately two-thirds of 48 living chickens showed diarrhea, 1 of 62 ducks showed lethargy, and 2 turkeys were asymptomatic. The household was located within a remnant fragment of the Dry Chaco biome, a hot and semi-arid tropical dry forest, surrounded by agriculture cropland. The affected flock had free access to a small pond (40 × 30 meters) frequently visited by wild waterfowl. The premise was depopulated and disinfected, and nearby backyard poultry within a 3 km perifocal zone (1 household) and 3–10 km surveillance zone (7 households) were inspected, with no evidence of morbidity or mortality. No affected wildlife were found in the visited premises.

Oropharyngeal and cloacal swabs samples from eight birds (seven chickens and one duck) were tested for influenza A virus and subtyped by real-time reverse transcription PCR (RRT-PCR) at SENASA reference laboratory. Positive samples underwent next-generation sequencing according to previously described protocols (6). Full and partial genome sequences were deposited in GISAID database (https://gisaid.org/) under accession codes EPI_ISL_19752381 and EPI_ISL_19823059–68.

Global avian influenza virus (AIV) phylogenies (including both HPAI and LPAI virus sequences) were inferred independently for each of the eight gene segments to determine the genetic composition of H5N1 viruses from this outbreak (hereafter referred as “H5N1-Arg_Feb2025 viruses”). The phylogenies indicate that the H5N1-Arg_Feb2025 viruses are novel 4:3:1 “triple reassortants” (**Figure 1**). Four segments (PB2, PB1, PA, and NS) belong to the South American LPAI lineage (shaded pink in **Figure 1**) that has been circulating in wild birds in the region for decades (10–13). Three segments (HA, NA, and MP) cluster with H5N1 viruses of the B3.2 genotype (**Figure 1**) and belong to the original Eurasian H5N1 lineage introduced to North America (**Figure 2**). One segment, NP, is exceptional. It does not cluster with any previously known South American viruses (HPAI or LPAI) and instead groups with North American LPAI viruses (**Figure 3**). Given limited surveillance in South America, where approximately 90% of full-genome sequences are from Argentina and Chile, it is difficult to determine how the North American-derived NP segment became part of the triple reassortant H5N1 here described.

**Figure 1.**
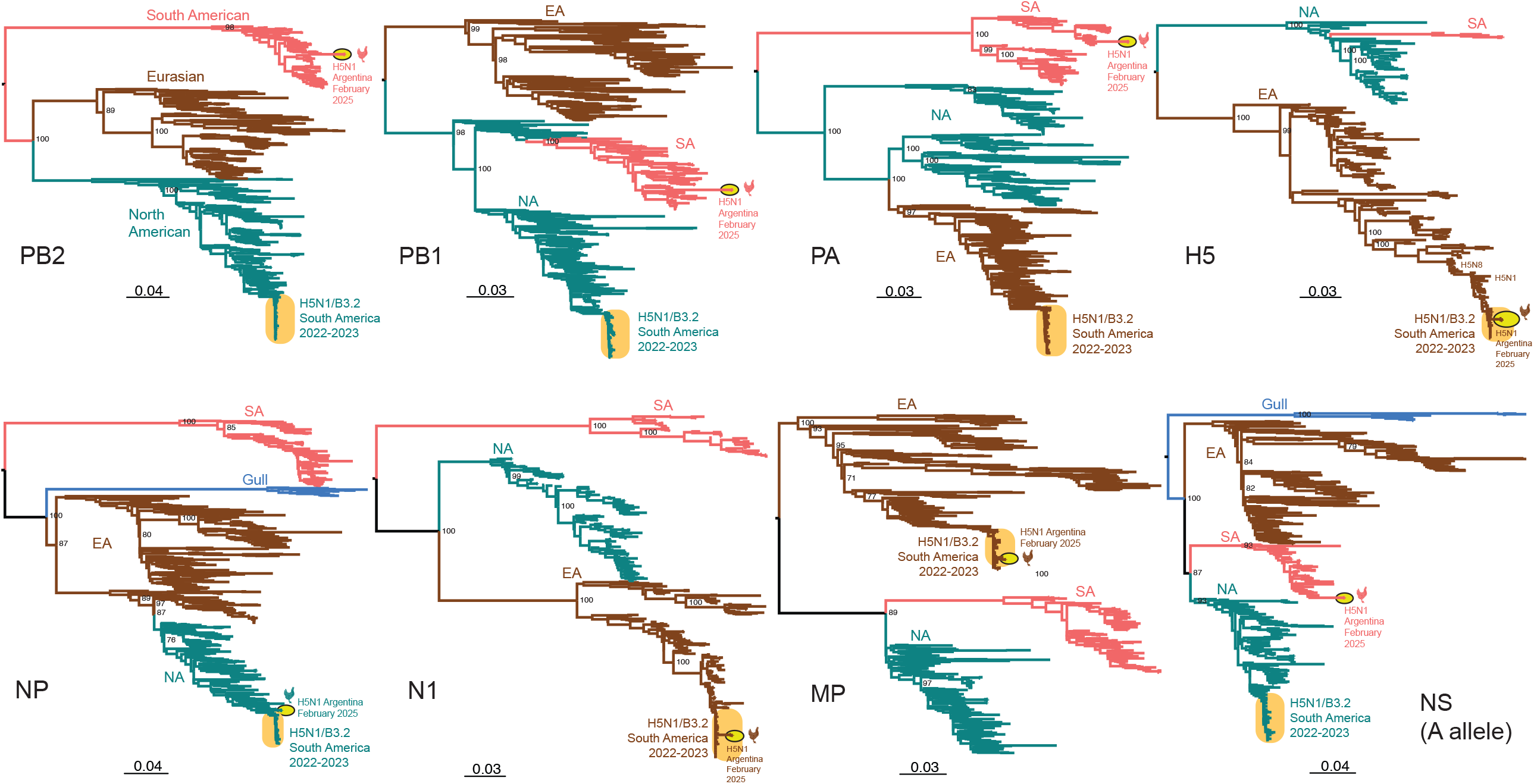
Maximum likelihood trees inferred for the eight IAV genome segments. Phylogenetic trees were inferred for each of the eight genome segments from IAVs collected globally in avian hosts (for NS only A allele is shown). Trees are midpoint rooted for clarity. Branch lengths are drawn to scale (nucleotide substitutions per site; scale-bars are shown). Bootstrap values are provided for key nodes. Branches are shaded by avian influenza lineage: North American (NA) lineage = teal, Eurasian (EA) lineage = brown, South American (SA) lineage = pink, Gull lineage (NP and NS segments) = blue. The previously reported South American H5N1 clade of B3.2 genotype viruses identified in multiple countries in avian and marine mammal hosts is highlighted in an orange oval and labeled. The new H5N1 viruses collected from backyard poultry in Argentina in February 2025 are highlighted in a yellow oval with black outline and labeled along with a chicken silhouette.

**Figure 2.**
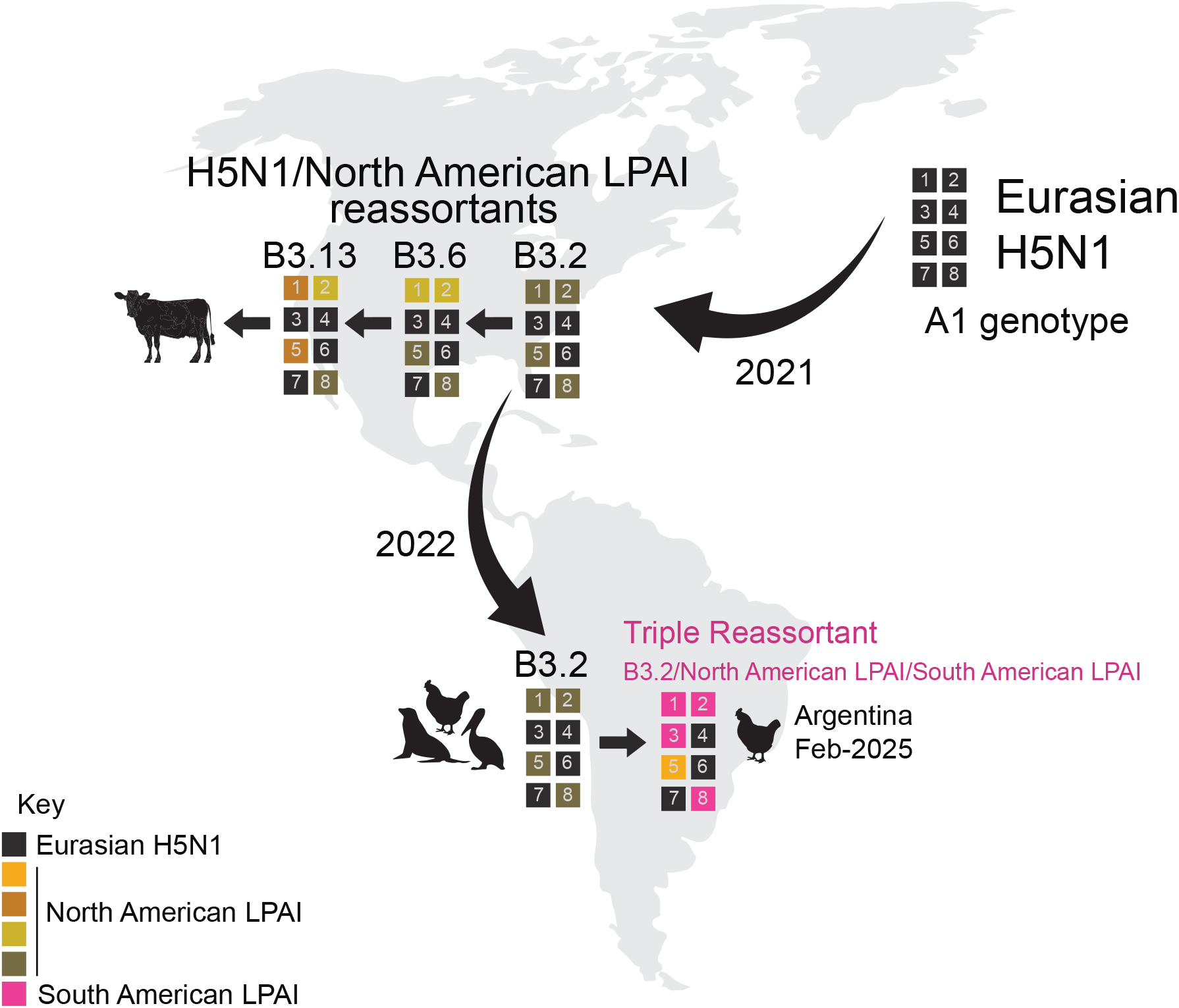
Key reassortment and migration events leading to the triple reassortant H5N1 viruses in February 2025, Argentina. Each box represents one of the eight segments of the IAV genome, numbered in order of longest to shortest length: 1-PB2, 2-PB1, 3-PA, 4-HA, 5-NP, 6-NA, 7-MP, and 8-NS. Each box is shaded according to the AIV lineage of that segment, with black = Eurasian H5N1, pink = South American LPAI, and various shades of orange/brown/yellow representing different clades of North American LPAI. Curved black arrows indicate the direction of major geographical migration events. Straight black arrows indicate sequential reassortment events of interest.

**Figure 3.**
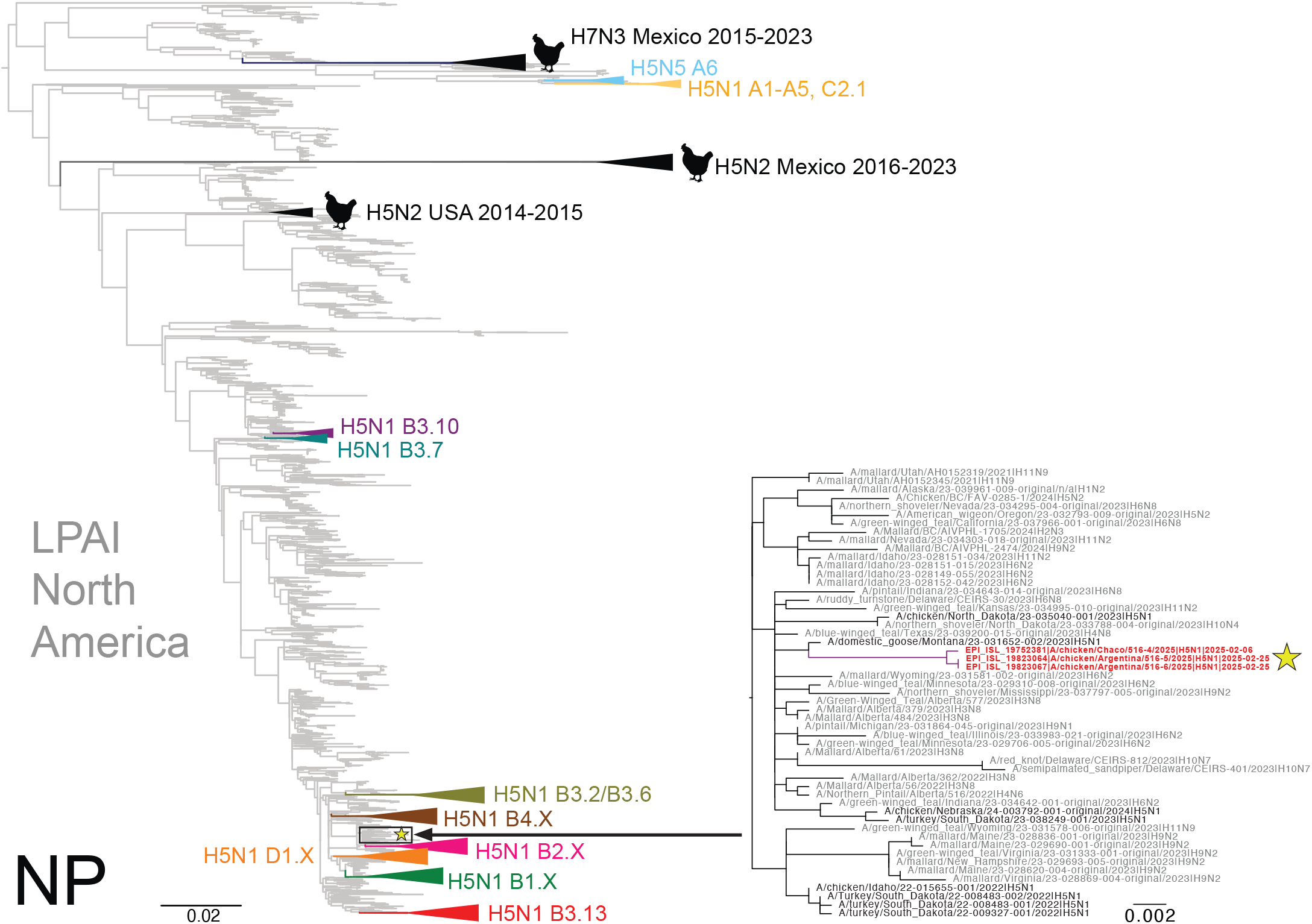
North American LPAI lineage contributes NP genes to H5N1 via reassortment. Phylogenetic tree inferred using ML methods for 11,820 NP sequences collected from LPAI and HPAI in the Americas, 2015–2025. LPAI is shaded grey. HPAI H5N1 clades are collapsed and shaded in different colors and labeled. Prior H5N2 and H7N3 outbreaks in poultry are shaded black. The three H5N1 viruses collected from poultry in Argentina in February 2025 are indicated in red and with a yellow star. A more detailed subsection of the tree containing these three viruses is provided with tip labels. Branch lengths are drawn to scale (nucleotide substitutions per site; scale-bar is shown).

To our knowledge, this represents the first documented reassortment event between HPAI H5N1 and endemic South American LPAI viruses. The South American PB2 and PA segments are highly divergent from the rest of global AIV diversity (10) (**Figure 1**), suggesting that reassortment involving these genes has significantly expanded the genetic diversity of H5N1 polymerase segments. Notably, although the H5N1-Arg_Feb2025 viruses have exchanged five gene segments, they retained the original Eurasian MP segment (**Figure 2**), which remains 100% conserved among H5N1 viruses circulating in North America. The absence of MP replacement by LPAI-derived segments, either in North or South America, suggests this MP segment may confer a selective advantage in HPAI H5 viruses. Currently, there is no evidence of this novel 4:3:1 triple reassortant in other South American countries; however, should future detections confirm wider spread, classification as a new H5N1 genotype would be warranted.

Reassortment is a key mechanism in the evolution and host adaptation of influenza viruses, often facilitating their emergence in new wildlife communities and contributing to the development of strains with panzootic or pandemic potential (14,15). The genomic constellation of the H5N1-Arg_Feb2025 viruses parallels patterns seen in North America, where clade 2.3.4.4b H5N1 viruses have also incorporated primarily internal LPAI genes. Particularly, the genome of South American novel H5N1 reassortant viruses resembles that of North American genotypes B3.6 and B3.13 (**Figure 2**), the latter widespread in U.S. dairy cattle, but with the Eurasian PA segment replaced by one from South American LPAI viruses. Despite substantial genomic changes, the novel Argentine reassortant viruses caused morbidity and mortality rates comparable to those previously observed for the B3.2 genotype, and available diagnostic techniques remained effective for their detection. However, the predominance of gastrointestinal rather than neurological signs suggests possible shifts in tissue tropism and/or virulence, warranting further investigation. Also, the detection of a North American NP segment not previously identified in LPAI viruses from Argentina or elsewhere in South America highlights the need to strengthen regional AIV surveillance, even in the absence of active HPAI circulation. There is a paucity of information on avian species and flyways involved in introductions of North American gene segments to South American LPAI viruses. Moreover, investigating the functional role of the North American-derived NP gene within the South American genomic background is needed to clarify its potential contribution to the distinct gastrointestinal phenotype observed in this outbreak.

In conclusion, our findings underscore the critical importance of sustained influenza surveillance coupled with whole-genome sequencing to track the evolution of HPAI H5N1 and support efforts to control and mitigate its impact on poultry, wildlife, and human health. Further research on the diversity of LPAI viruses circulating in Neotropical wildlife will be essential to understand potential interactions between H5N1 and South American lineage strains, and to assess the long-term consequences of the introduction of HPAI viruses into the region (and potentially beyond, should these reassortant strains spread to other regions).

## Contributions

R.E.V.T, M.U., and A.R. conceptualized the research. A.P. was responsible for logistics to collect samples during the outbreak and submission to the laboratory. M.C.A., V.D.M., L.E.P., and E.P. performed molecular diagnosis, subtypification and genome sequencing. M.I.N. and A.C.B. performed phylogenies. R.E.V.T, M.I.N., and A.R. wrote and prepared the original manuscript draft. All authors reviewed and edited the manuscript and agreed to the final version.

## ACKNOWLEDGMENTS

This work was conducted under a specific agreement between the National Administration of Laboratories and Health Institutes (ANLIS) and the National Service of Agri-food Health and Quality (SENASA), Argentina.

